# Amino Acid-Stimulated Muscle Protein Synthesis is Attenuated During Early Recovery from Aerobic Exercise in Individuals with Obesity

**DOI:** 10.64898/2026.06.14.732200

**Authors:** Kailin A. Johnsson, Eduardo D. S. Freitas, Lori R. Roust, Eleanna De Filippis, Lingjun Li, Haiwei Gu, Matthew R. Buras, Christos S. Katsanos

## Abstract

**Background:** In healthy individuals, exercise and amino acid availability act synergistically to stimulate muscle anabolism. However, this interaction may be impaired in individuals with obesity.

**Objective:** We examined whether acute aerobic exercise alters amino acid-stimulated muscle protein synthesis during immediate postexercise recovery in adults with obesity.

**Methods:** Sixteen sedentary adults with a body mass index >30 kg/m^2^ underwent a stable-isotope tracer infusion to measure mixed-muscle protein fractional synthesis rate (FSR) in the basal/fasted state and during an amino acid infusion, either with or without prior aerobic exercise. Participants were randomly assigned to receive either amino acid infusion alone (AA) or 45 min of cycling exercise at ∼65% heart rate reserve immediately before amino acid infusion (EX + AA).

**Results:** Amino acid infusion increased muscle protein FSR in the AA group (*P* < 0.0001) but not in the EX + AA group (*P* > 0.05), and the amino acid-stimulated increase in FSR was 78% lower in EX + AA than in AA (*P* < 0.01). The amino acid infusion increased (*P* < 0.05) plasma amino acid concentrations in both groups. However, during the amino acid infusion, plasma concentrations of essential and branched-chain amino acids, including leucine, were lower in EX + AA than in AA (*P* < 0.05). Moreover, across participants, absolute changes in muscle protein FSR were positively associated with plasma leucine concentrations during the amino acid infusion (*P* < 0.05).

**Conclusions:** These findings show that, in individuals with obesity, acute aerobic exercise markedly attenuates amino acid-stimulated muscle protein synthesis during early postexercise recovery. These findings have important implications for designing nutritional strategies to optimize muscle anabolism in this population.

## INTRODUCTION

Although obesity has traditionally been viewed primarily as a disorder characterized by impairments in skeletal muscle glucose and lipid metabolism (1, 2), growing evidence indicates that obesity also disrupts skeletal muscle protein metabolism (3, 4). Increased circulating amino acid availability is a principal nutritional stimulus for muscle protein synthesis (5, 6), and attenuated muscle protein synthetic responses to amino acid provision have been reported in individuals with obesity (7), suggesting the presence of obesity-associated anabolic resistance.

Exercise is a potent regulator of skeletal muscle protein metabolism. In healthy individuals, exercise and increased amino acid availability interact to amplify the stimulation of muscle protein synthesis (8, 9). This effect has been studied primarily in the context of resistance exercise, which potentiates the responsiveness of skeletal muscle to increased amino acid availability (10). The interaction between exercise and nutrient provision is considered a fundamental determinant of skeletal muscle adaptation, and forms the physiological basis for recommendations advocating the coupling of exercise with adequate dietary protein intake (11). However, emerging evidence suggests that obesity may impair this interaction. For example, Beals et al. (12) showed that obesity attenuated the capacity of prior resistance exercise to enhance fed-state muscle protein synthesis. While resistance exercise has received the greatest attention in the context of nutrient-stimulated muscle protein synthesis, aerobic exercise also has the capacity to increase muscle protein synthesis (13), albeit to a lesser magnitude than resistance exercise when performed immediately before nutrient provision (14). Understanding the interaction between aerobic exercise and plasma amino acid availability specifically in individuals with obesity is particularly important because aerobic exercise is among the most widely prescribed interventions for obesity management. Nevertheless, it remains unclear whether aerobic exercise potentiates or attenuates amino acid-stimulated muscle protein synthesis in this population.

Aerobic exercise does not appear to alter muscle protein synthesis during the exercise session itself (15), but protein synthesis increases during the postexercise recovery period in healthy adults (13, 16). In the postabsorptive state in healthy humans, aerobic exercise has been reported to increase mixed-muscle protein synthesis rates by approximately 45% (15), with comparable increases reported in subsequent studies (17). However, the immediate postexercise period is characterized by a transient, exercise-induced cellular stress in muscle (18, 19), which appears to be exacerbated by obesity (20). This altered postexercise environment could therefore limit skeletal muscle responsiveness to increased plasma amino acid availability, thereby causing aerobic exercise to transiently attenuate, rather than enhance, the anabolic response to amino acids in individuals with obesity. This possibility is particularly relevant given that hyperaminoacidemia alone effectively stimulates muscle protein synthesis (5), including in individuals with obesity (21). Therefore, the aim of this study was to determine whether acute aerobic exercise alters amino acid-stimulated muscle protein synthesis during early postexercise recovery in individuals with obesity. We hypothesized that prior aerobic exercise would attenuate the muscle protein synthetic response to increased amino acid availability during this period.

## METHODS

### Study Participants

We studied sixteen individuals with obesity (body mass index [BMI] >30 kg/m^2^) who did not engage in regular physical activity or exercise more than two days per week and did not meet current physical activity recommendations (22). All study procedures were approved by the Institutional Review Board at Mayo Clinic, and written informed consent was obtained from each participant. Prior to obtaining written consent, the study purpose, requirements, and potential risks associated with study participation were explained to each participant. The studies from which these data were derived were registered at ClinicalTrials.gov as NCT01824173 and NCT04700800.

Participants first underwent screening within the Ambulatory Infusion Center (AIC) at Mayo Clinic in Arizona. Eligibility was determined based on medical history, physical examination, electrocardiogram, and standard blood and urine analyses, confirming that all participants were otherwise healthy. Exclusion criteria included the presence of acute illness, diabetes, liver, renal, or cardiovascular disease, and chronically elevated blood pressure (systolic, >150 mmHg; diastolic, >100 mmHg). Participants were also excluded if they were actively engaged in a weight-loss program, smoked, or used nutritional supplements, prescription medications, or over-the-counter drugs. During the screening visit, insulin sensitivity was estimated using the Matsuda-Insulin Sensitivity Index (ISI) (23). The Matsuda-ISI was calculated from plasma insulin and glucose responses obtained during an oral glucose tolerance test (OGTT), as previously described (23). Following the OGTT, participants were included only if fasting plasma glucose was <126 mg/dL and 2-h plasma glucose was <200 mg/dL, consistent with the absence of diabetes.

Participants returned to the AIC on a separate day for assessment of body composition using dual-energy X-ray absorptiometry (Hologic, Inc). This assessment was followed by the determination of peak oxygen consumption (VO_2_peak) using an incremental cycle ergometer test. For this test, after a 5-min unloaded warm-up, work rate was increased by 20 W·min^-1^ while participants were instructed to maintain a pedaling cadence of 65 revolutions·min^-1^ until volitional exhaustion. During the VO_2_peak test, a 12-lead electrocardiogram was continuously recorded, and blood pressure and oxygen saturation were monitored throughout the test. Participants were verbally encouraged to exercise until exhaustion (24), and all achieved a respiratory exchange ratio >1.1 prior to test termination, consistent with maximal effort (25). Expired gases were continuously analyzed using a metabolic cart (MedGraphics, Saint Paul, MN).

### Experimental Design

On a separate day, participants arrived at the AIC at ∼7:00 AM following a ∼10-h overnight fast. A stable-isotope tracer infusion was used to measure skeletal muscle protein synthesis in the basal/fasted state and during an intravenous amino acid infusion designed to increase plasma amino acid concentrations to levels previously shown by our group to stimulate muscle protein synthesis, including in individuals with obesity (21). Eight participants assigned to the exercise group performed a single session of aerobic exercise after completion of basal measurements and immediately before initiation of the amino acid infusion (EX + AA group). Another eight participants received the amino acid infusion alone (AA group) and served as the control group. Participants were randomly assigned to each group in a parallel-design trial. All participants were instructed to refrain from structured exercise, maintain their habitual diet, and avoid alcohol consumption for the 3 days preceding each study visit. Adherence to fasting and physical activity instructions was confirmed prior to initiation of the tracer infusion study.

While resting in bed, participants had one catheter placed into an antecubital arm vein for infusion of d_10_ leucine (L-[2,3,3,4,5,5,5,6,6,6-^2^H_10_]leucine), at a rate of 0.15 μmol·kg FFM^- 1^·min^-1^ (priming dose, 6.4 μmol·kg FFM^-1^), and which was maintained throughout the study to assess mixed-muscle protein synthesis rates. A separate catheter was placed retrogradely in a dorsal hand vein for blood sampling using the heated-hand technique. Following collection of samples associated with the basal state, a primed continuous infusion of an amino acid solution (15% Clinisol, Baxter Healthcare Corporation, Deerfield, IL) was initiated at 4 mg·kg FFM^-1^·min^-1^ (priming dose, 82 mg·kg FFM^-1^) and continued for 240 mins. During the amino acid infusion, the infusion rate of d_10_-leucine was increased to 0.29 µmol·kg FFM^-1^·min^-1^ (priming dose, 2.6 µmol·kg FFM^-1^) to account for tracer dilution resulting from the exogenous (i.e., infused) leucine delivery. For the EX + AA group, participants performed 45 min of cycling exercise prior to the initiation of the amino acid infusion, which was administered for the same duration as in the AA group (i.e., 240 min) immediately following completion of the exercise session. The exercise was performed at an intensity corresponding to 65% of heart rate reserve, calculated from each participant’s resting heart rate and the maximal heart rate achieved during the VO_2_peak test (26). During the exercise session, the workload on the cycle ergometer was adjusted as needed to maintain heart rate within ±5 bpm of the target exercise intensity. Following completion of the exercise session, participants rested in bed for the remainder of the study, and all subsequent procedures were identical to those performed in the AA group. The overall experimental design is depicted in **Figure 1**.

**Figure 1.**
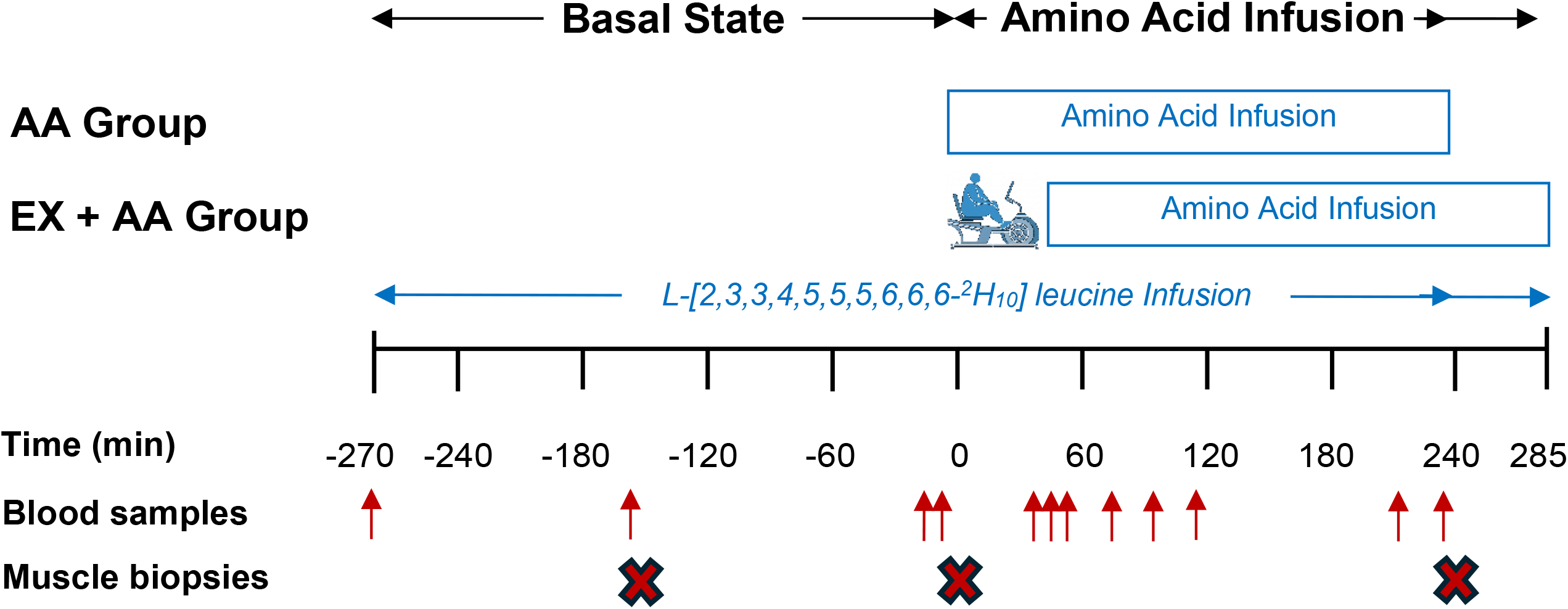
Experimental design. Participants underwent a continuous leucine tracer infusion during Basal State and Amino Acid Infusion periods. One group received the amino acid infusion without prior exercise (AA), whereas the other group performed 45 min of aerobic exercise immediately before initiation of the amino acid infusion (EX + AA). Blood samples and muscle biopsies were collected at the indicated time points. For clarity, sampling time points during the Amino Acid Infusion period are shown only for the AA group. The corresponding sampling time points for the EX + AA group occurred 45 min later than those for the AA group, following completion of the exercise session.

Two muscle biopsy samples (∼70 mg each) were collected from the vastus lateralis muscle during the basal state period. A third biopsy was collected at the end of each group’s respective amino acid infusion period, which occurred 45 min later in EX + AA because of the preceding exercise session (Figure 1). Muscle tissue samples were immediately cleaned of visible fat and connective tissue, blotted dry, and frozen in liquid nitrogen. Blood samples were collected at selected time points during the basal and the corresponding amino acid infusion periods (Figure 1) to determine d_9_-leucine enrichment as we have previously described (27) and to measure selected biochemical parameters.

### Sample Analyses

Leucine enrichment (i.e., d_9_-leucine) was measured in blood and mixed-muscle protein using procedures we have previously described (21, 27). Briefly, proteins in 1 mL of blood were precipitated using sulfosalicylic acid (SSA). Mixed-muscle samples (∼15 mg) were similarly processed by homogenization in SSA to precipitate proteins. Proteins in both blood and mixed-muscle samples were subsequently hydrolyzed with 6 N HCl. Amino acids were then isolated using cation-exchange chromatography, and d_9_-leucine enrichment was determined by liquid chromatography-tandem mass spectrometry (LC-MS/MS) at the Mayo Clinic Metabolomics Core. Blood d_9_-leucine enrichment and mixed-muscle protein-bound d_9_-leucine enrichments were used to calculate the mixed-muscle protein fractional synthesis rates (FSR; %·h^-1^) during the Basal period (between biopsies 1 and 2), and the Amino Acid Infusion period (between biopsies 2 and 3) using the precursor-product approach, as previously described (27).

Plasma amino acid concentrations were measured by LC-MS/MS using a method adapted from previously published protocols (28–30). Analyses were performed using an Agilent 1290 UPLC–6495 QQQ-MS (Santa Clara, CA) system, with chromatographic separation conducted under hydrophilic interaction chromatography groups on a Waters XBridge BEH Amide column (Waters Corporation, Milford, MA). The mass spectrometer was operated with an electrospray ionization source, and amino acids were quantified using chemical standards under the targeted multiple-reaction monitoring mode. Peak integration was performed with Agilent MassHunter Quantitative Data Analysis software (Santa Clara, CA). Individual plasma amino acid concentrations were quantified at 10 min before the end of the basal period and at 55 and 210 min after initiation of the amino acid infusion (Figure 1). Plasma amino acid concentrations were also reported as the sum of all measured amino acids (TAA), essential amino acids (EAA), branched-chain amino acids (BCAA), and non-essential amino acids (NEAA), and given the central role of EAA, and particularly BCAA, in stimulating skeletal muscle protein synthesis (5, 31, 32).

Plasma glucose and insulin concentrations were measured at 10 min before the end of the basal period and at 40, 55, 70, 90, 110, and 210 min after initiation of the amino acid infusion. Plasma glucose concentrations were determined using an automated glucose analyzer (YSI 2300, Yellow Springs, OH). Plasma insulin concentrations were determined using a commercially available enzyme-linked immunosorbent assay kit (80-INSHU-E01.1; ALPCO Diagnostics). Clinical chemistry parameters assessed during the screening visit were analyzed at the Mayo Clinic Laboratories (Mayo Clinic, Arizona).

### Statistical Analyses

Data normality was determined using the Levene’s test and supplemented by graphical information. Baseline participant characteristics (**Table 1**) were compared between groups using unpaired Student’s *t*-tests. A two-way repeated-measures ANOVA [group (AA and EX + AA) x time (before and during the amino acid infusion)] was used to assess the main effects of group and time and the group × time interaction for the primary outcome (i.e., muscle protein FSR) and secondary outcomes (i.e., plasma amino acid concentrations, insulin, glucose). Pairwise comparisons were conducted using Bonferroni adjustment for multiple comparisons. Changes in muscle protein FSR from basal were compared between groups using an unpaired Student’s t-test. Associations between variables of interest were evaluated using Pearson’s product-moment correlation coefficient (*r*). Data are presented as mean ± SEM, and statistical significance was set at *P* < 0.05. All statistical analyses were performed using GraphPad Prism (Version 10; GraphPad Software, Boston, MA).

**Table 1.**
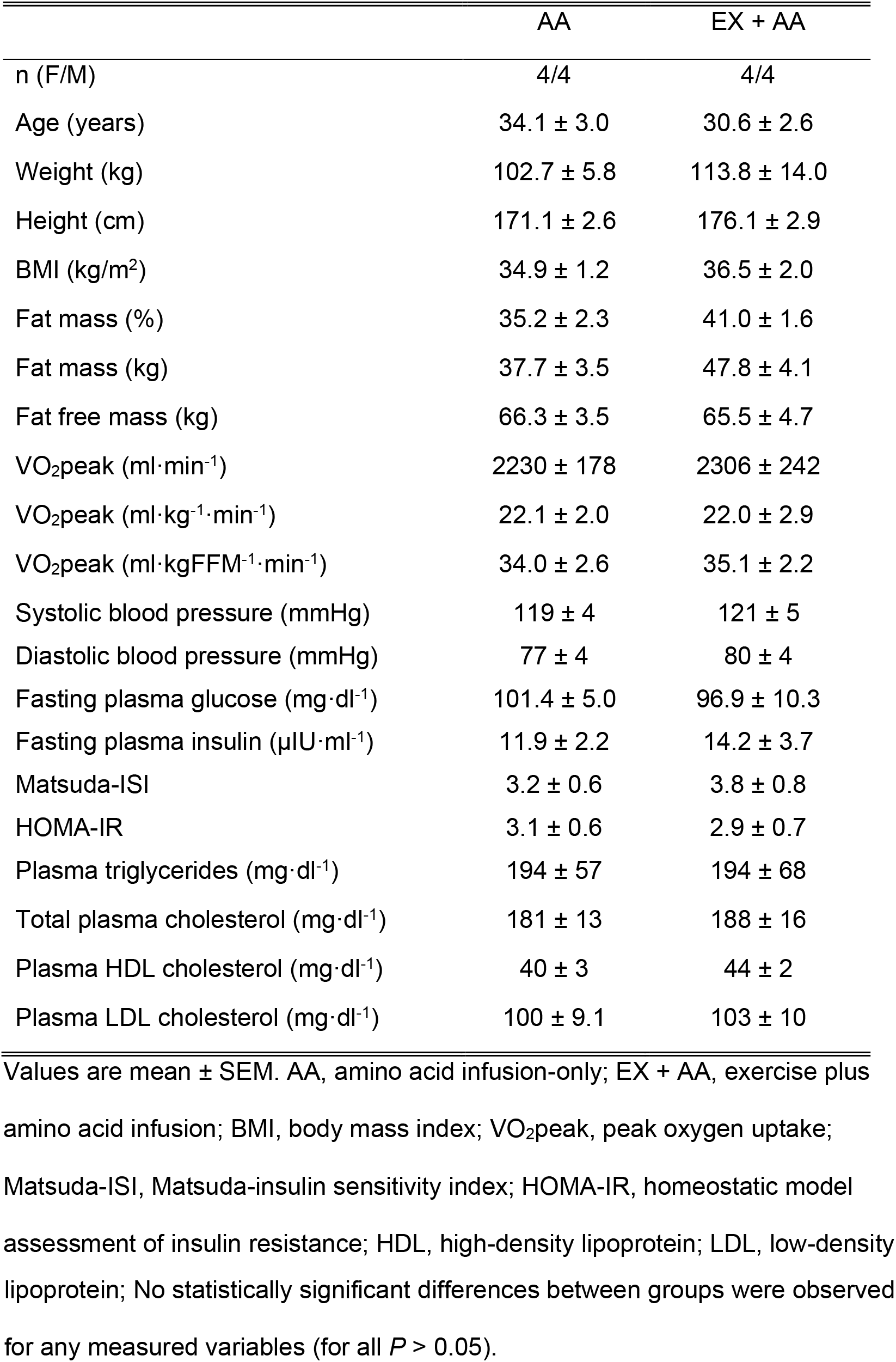
Participant characteristics.

## RESULTS

Anthropometric characteristics and fasting blood chemistry parameters did not differ significantly between groups (**Table 1**). Consistent with the physical inactivity criterion applied during the screening, mean VO_2_peak was ∼22 ml·kg^-1^·min^-1^ in both groups, indicative of low cardiorespiratory fitness according to established population reference standards (33).

### Muscle Protein Synthesis

Muscle protein synthesis, expressed as FSR and assessed as the integrated response across exercise and recovery was the primary endpoint of the study. Blood d_9_-leucine enrichment responses during the Basal and the Amino Acid Infusion periods in both groups are shown in **Figure 2**. No significant main effects of group or amino acid infusion, and no group x amino acid infusion interaction, were observed for plasma d_9_-leucine enrichment (for all *P >* 0.05). For muscle protein synthesis, two-way ANOVA showed a significant main effect of amino acid infusion on muscle protein synthesis rate (*P* < 0.001), no significant main effect of group (*P* >0.05), and a significant group x amino acid infusion interaction (*P* < 0.01). Basal muscle protein FSR did not differ between groups (*P* > 0.05). Pairwise comparisons showed that the amino acid infusion significantly increased mixed-muscle protein FSR from the basal period in the AA group (*P* < 0.0001), whereas FSR did not increase significantly in the EX + AA group (*P >* 0.05; **Figure 3A**). Figure 3A also depicts individual participants’ FSR responses to the amino acid infusion. The corresponding absolute changes in FSR are summarized in **Figure 3B**, showing that the amino acid-stimulated increase in FSR was 78% lower in the EX + AA group than in the AA group (95% CI: −0.0761 to −0.0158; *P* < 0.01).

**Figure 2.**
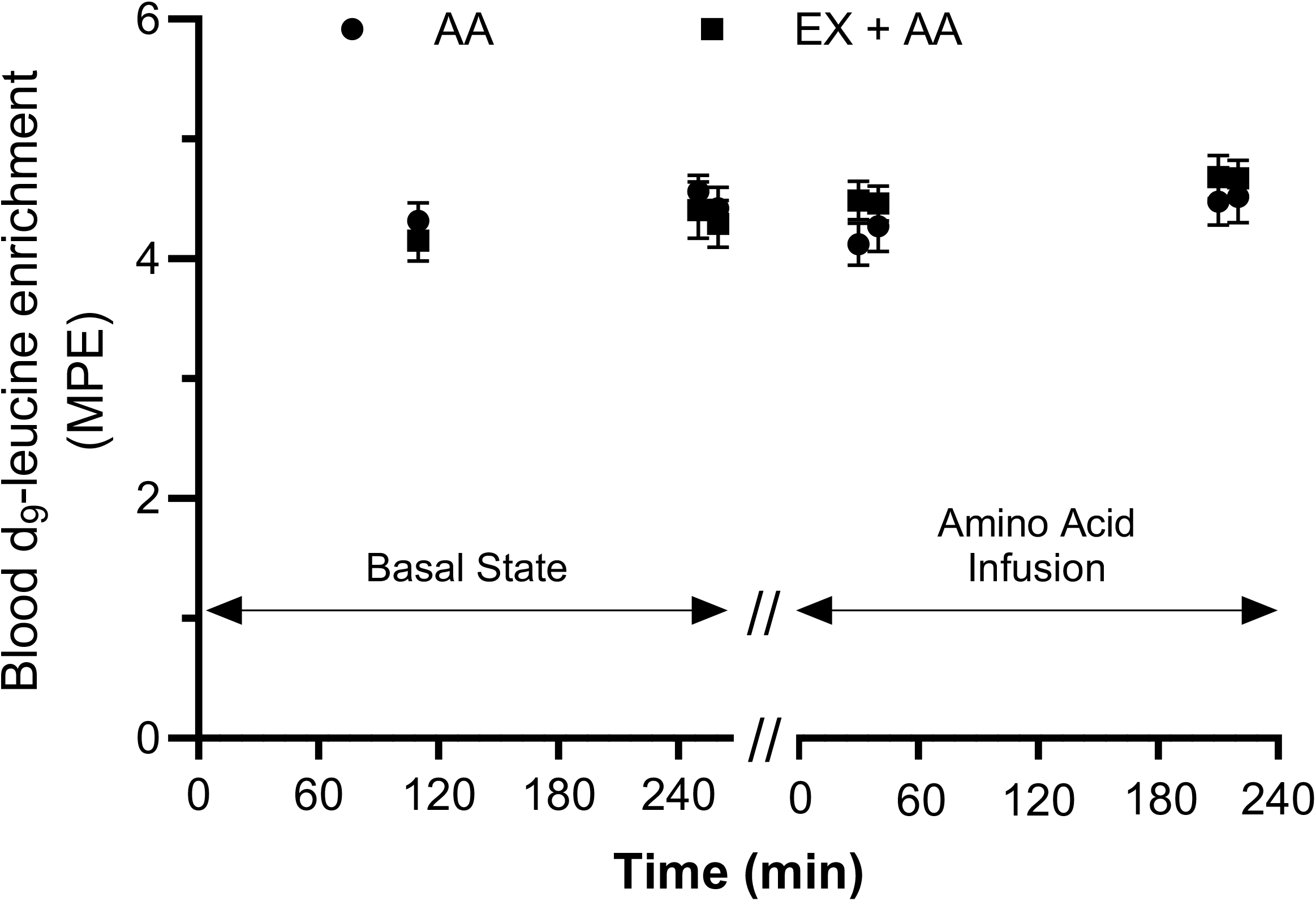
Blood d_9_-leucine enrichment. Blood d_9_-leucine enrichment is shown during the Basal State and Amino Acid Infusion periods in the amino acid infusion-only (AA) and exercise plus amino acid infusion (EX + AA) groups. Values are presented as means ± SEM. N = 8 per group. The break in the x-axis indicates the transition between the basal and amino acid infusion periods.

**Figure 3.**
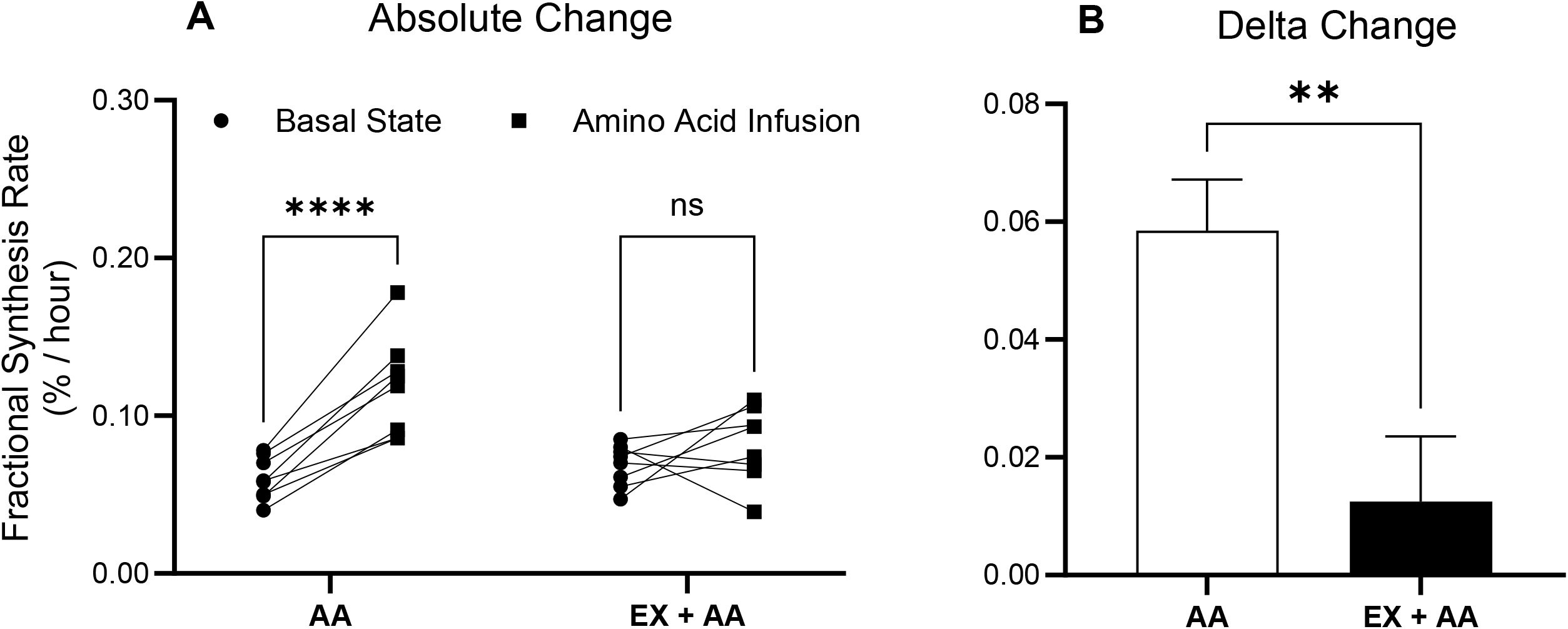
Fractional synthesis rate of mixed-muscle protein. Mixed-muscle protein fractional synthesis rate (FSR) was measured during the Basal State and Amino Acid Infusion periods in the amino acid infusion-only (AA) and exercise plus amino acid infusion (EX + AA) groups. (A) Individual participant muscle protein FSR responses during the Basal State and Amino Acid Infusion periods. (B) Absolute change in muscle protein FSR from the Basal State to the Amino Acid Infusion period. Values in panel B are presented as means ± SEM. N = 8 per group. ****, *P* < 0.0001; **, *P* < 0.01; ns, not statistically significant.

### Plasma Amino Acids

**Table 2** shows plasma concentrations of individual amino acids, as well as TAA, EAA, BCAA, and NEAA, during the Basal (i.e., 10 mins prior to the end of the Basal Period) and the Amino Acid Infusion (i.e., average of the 55 and 210 mins after initiation of the amino acid infusion) periods. ANOVA revealed a significant main effect of amino acid infusion (time; *P* < 0.05) in the concentration for all measured plasma amino acids, with the exception of tyrosine and asparagine, as well as for all grouped amino acid categories. Significant main effects of group (*P* < 0.05) were observed only for plasma arginine, leucine, and phenylalanine concentrations. Table 2 also shows the corresponding delta changes from basal in the plasma amino acid concentrations during the amino acid infusion within the AA and EX + AA groups. These responses differed between groups (*P* < 0.05) for plasma alanine, arginine, leucine, methionine, phenylalanine, serine, threonine, and valine. Specifically, the increase was lower in the EX + AA group for plasma alanine, leucine, methionine, phenylalanine, serine, and valine, but higher for arginine and threonine concentrations. Accordingly, the increases in plasma TAA, EAA, and BCAA, but not NEAA, concentrations were significantly lower (*P* < 0.05) in the EX + AA group compared with the AA group during the amino acid infusion.

**Table 2.** Plasma amino acid concentrations during the Basal State and Amino Acid Infusion periods, and corresponding absolute changes from Basal State.

| | Basal State | Amino Acid<br>Infusion | Change ($\Delta$) |
| --- | --- | --- | --- |
| Alanine |  |  |  |
| AA | 311 ± 37 | 598 ± 51<sup>***</sup> | 287 ± 21 |
| EX + AA | 335 ± 30 | 543 ± 29<sup>***</sup> | 187 ± 17<sup>††</sup> |
| Arginine |  |  |  |
| AA | 59 ± 5 | 198 ± 4<sup>***</sup> | 139 ± 5 |
| EX + AA | 76 ± 11 | 277 ± 16<sup>***</sup> | 202 ± 9<sup>†††</sup> |
| Asparagine |  |  |  |
| AA | 27 ± 2 | 24 ± 2 | -3 ± 3 |
| EX + AA | 32 ± 2 | 24 ± 1<sup>*</sup> | -7 ± 3 |
| Glutamic Acid |  |  |  |
| AA | 47 ± 6 | 97 ± 13<sup>**</sup> | 50 ± 8 |
| EX + AA | 53 ± 6 | 125 ± 14<sup>***</sup> | 72 ± 12 |
| Glycine |  |  |  |
| AA | 290 ± 22 | 551 ± 27<sup>***</sup> | 262 ± 13 |
| EX + AA | 310 ± 37 | 566 ± 40<sup>***</sup> | 256 ± 21 |
| Histidine |  |  |  |
| AA | 68 ± 6 | 117 ± 7<sup>***</sup> | 50 ± 12 |
| EX + AA | 77 ± 8 | 122 ± 6<sup>***</sup> | 46 ± 7 |
| Isoleucine |  |  |  |
| AA | 66 ± 4 | 205 ± 7<sup>***</sup> | 139 ± 7 |
| EX + AA | 69 ± 7 | 208 ± 9<sup>***</sup> | 140 ± 3 |
| Leucine |  |  |  |
| AA | 137 ± 8 | 431 ± 16<sup>***</sup> | 294 ± 16 |
| EX + AA | 139 ± 9 | 213 ± 14<sup>***</sup> | 74 ± 7<sup>†††</sup> |
| Methionine |  |  |  |
| AA | 46 ± 1 | 398 ± 24<sup>***</sup> | 352 ± 23 |
| EX + AA | 56 ± 6 | 317 ± 23<sup>***</sup> | 261 ± 19<sup>††</sup> |
| Phenylalanine |  |  |  |
| AA | 62 ± 2 | 261 ± 7<sup>***</sup> | 199 ± 6 |
| EX + AA | 71 ± 5 | 184 ± 10<sup>***</sup> | 113 ± 5<sup>†††</sup> |
| Serine |  |  |  |
| AA | 93 ± 2 | 174 ± 4<sup>***</sup> | 81 ± 5 |
| EX + AA | 104 ± 7 | 170 ± 8<sup>***</sup> | 66 ± 4<sup>†</sup> |
| Threonine |  |  |  |
| AA | 125 ± 14 | 236 ± 17<sup>***</sup> | 111 ± 6 |
| EX + AA | 143 ± 25 | 300 ± 31<sup>***</sup> | 156 ± 15<sup>†</sup> |
| Tyrosine |  |  |  |
| AA | 75 ± 1 | 80 ± 1 | 5 ± 0 |
| EX + AA | 83 ± 11 | 89 ± 11 | 6 ± 8 |
| Valine |  |  |  |
| AA | 226 ± 7 | 558 ± 34<sup>***</sup> | 332 ± 29 |
| EX + AA | 245 ± 20 | 462 ± 22<sup>***</sup> | 217 ± 17<sup>††</sup> |
| Total Amino Acids |  |  |  |
| AA | 1634 ± 55 | 3932 ± 148<sup>***</sup> | 2297 ± 108 |
| EX + AA | 1814 ± 117 | 3598 ± 152<sup>***</sup> | 1788 ± 86<sup>††</sup> |
| Essential Amino Acids |  |  |  |
| AA | 731 ± 13 | 2207 ± 82<sup>***</sup> | 1476 ± 79 |
| EX + AA | 801 ± 62 | 1807 ± 82<sup>***</sup> | 1005 ± 42<sup>†††</sup> |
| Branched Chain Amino Acids |  |  |  |
| AA | 430 ± 14 | 1194 ± 51<sup>***</sup> | 765 ± 49 |
| EX + AA | 453 ± 30 | 883 ± 36<sup>***</sup> | 430 ± 23<sup>†††</sup> |
| Non-essential Amino Acids |  |  |  |
| AA | 903 ± 55 | 1724 ± 75<sup>***</sup> | 821 ± 35 |
| EX + AA | 1013 ± 62 | 1791 ± 78<sup>***</sup> | 783 ± 47 |

Values are in $\mu\text{mol}\cdot\text{L}^{-1}$ ; Data are mean ± SEM; AA, amino acid infusion only; EX + AA, exercise plus amino acid infusion; \*\*\*, $P < 0.001$ ; \*\*, $P < 0.01$ ; \*, $P < 0.05$ versus Basal; †††, $P < 0.001$ ; ††, $P < 0.01$ ; †, $P < 0.05$ versus AA.

Correlation analyses across study participants revealed that the amino acid-stimulated change in muscle protein FSR from basal was associated with the plasma concentrations of specific amino acids measured during the amino acid infusion. Specifically, the change in muscle protein FSR was inversely correlated with plasma arginine (*r* = −0.62; 95% CI: −0.8541 to −0.1817; *P* = 0.01) and glycine (*r* = −0.50; 95% CI: −0.8000 to −0.01138; *P* = 0.04) concentrations and positively correlated with plasma leucine concentration (*r* = 0.62; 95% CI: 0.1726 to 0.8515; *P* = 0.01). However, the change in muscle protein FSR was not significantly correlated with the plasma concentrations of the other branched-chain amino acids, isoleucine (*P* = 0.93) or valine (*P* = 0.43). Furthermore, the change in muscle protein FSR was not significantly correlated with the plasma concentrations of TAA (*P* = 0.97), EAA (*P* = 0.28), BCAA (*P* = 0.07), or NEAA (*P* = 0.15) during the amino acid infusion.

### Plasma Glucose and Insulin

**Table 3** shows plasma glucose and insulin concentrations averaged across measurements obtained during the Basal (i.e., 10 mins prior to the end of the Basal Period) and the Amino Acid Infusion (i.e., average of 40, 55, 70, 90, 110, and 210 min after initiation of the amino acid infusion) periods, together with the corresponding absolute changes from Basal values. ANOVA revealed a significant main effect of group (*P* < 0.01), but not time, for plasma glucose concentrations, whereas a significant main effect of time (*P* < 0.01), but not group, was observed for plasma insulin concentrations. Amino acid infusion significantly increased plasma insulin concentrations in both the AA and EX + AA groups (Table 3). During the amino acid infusion, mean changes from basal in plasma glucose and insulin concentrations were 83% and 34% lower, respectively, in the EX + AA group than in the AA group (Table 3); however, neither between-group difference reached statistical significance (*P* > 0.05).

**Table 3.** Plasma glucose and insulin concentrations during the Basal State and Amino Acid Infusion periods, and corresponding absolute changes from Basal State.

| | Basal State | Amino Acid<br>Infusion | Change ($\Delta$) |
| --- | --- | --- | --- |
| Glucose |  |  |  |
| AA | 93.1 ± 3.9 | 96.7 ± 3.4 | 3.6 ± 2.9 |
| EX + AA | 84.3 ± 2.6 | 84.9 ± 2.2 | 0.6 ± 1.5 |
| Insulin |  |  |  |
| AA | 18.7 ± 3.4 | 48.3 ± 4.7*** | 29.7 ± 5.5 |
| EX + AA | 19.9 ± 3.2 | 39.4 ± 6.5** | 19.5 ± 4.7 |

Values are in mg·dL<sup>-1</sup> (glucose) and $\mu$ U·mL<sup>-1</sup> (insulin); Data are mean ± SEM; AA, amino acid infusion only; EX + AA, exercise plus amino acid infusion; \*\*\*, $P < 0.001$ ; \*\*, $P < 0.01$ versus Basal.

## DISCUSSION

The primary finding of the present study is that, in individuals with obesity, acute aerobic exercise markedly attenuated the amino acid-stimulated increase in muscle protein synthesis assessed as the integrated response across exercise and recovery period. Although amino acid infusion alone robustly stimulated muscle protein synthesis, no statistically significant increase above basal was detected when the amino acid infusion followed exercise, and the magnitude of this anabolic response was approximately 80% lower than in the amino acid infusion alone group. Together, these findings indicate substantially impaired muscle anabolic responsiveness to increased amino acid provision in obesity during the immediate recovery period from aerobic exercise.

In lean, healthy individuals, muscle protein synthesis remains unchanged during aerobic exercise itself (15). Consistent with this observation, under conditions comparable to those used in the present study, we previously found no change from basal in muscle protein synthesis measured as the integrated response across exercise and recovery in lean individuals and individuals with obesity (34). These findings suggest that, under the present similar experimental conditions, aerobic exercise alone is not expected to modify muscle protein synthesis. By contrast, when aerobic exercise is combined with increased plasma amino acid availability in lean individuals, muscle protein anabolism is enhanced during and immediately after exercise (35–37). Collectively, these studies of combined exercise and amino acid provision support the concept that aerobic exercise enhances the anabolic response of skeletal muscle to amino acid availability (37). To our knowledge, the present study is the first to examine this interaction specifically in individuals with obesity. This question is of particular clinical relevance because aerobic exercise is commonly prescribed to increase energy expenditure and improve glycemic control and cardiometabolic health. The divergent responses observed in the AA and EX + AA groups indicate that increased amino acid availability remains capable of stimulating muscle protein synthesis in individuals with obesity under nonexercise conditions, but that the anabolic response is markedly diminished when amino acid provision immediately follows aerobic exercise. Thus, our findings suggest that the enhanced amino acid responsiveness reported following aerobic exercise in lean individuals (35–37) may not extend to individuals with obesity.

Consistent with the robust response we observed in the AA group, increased plasma amino acid availability has previously been shown to stimulate muscle protein synthesis in individuals with obesity (21, 38, 39). Nevertheless, preserved responsiveness under some experimental conditions does not preclude obesity-associated anabolic resistance under others. Previous studies have demonstrated that obesity can reduce the responsiveness of skeletal muscle protein synthesis to anabolic stimuli such as insulin and feeding (7, 40). Relevant to the present findings, the muscle protein synthetic response to feeding immediately after resistance exercise is attenuated in individuals with obesity (12). The present study extends these observations to aerobic exercise. In the previous resistance-exercise study, feeding produced a reduced but still significant increase in muscle protein synthesis (12). In contrast, amino acid provision did not produce a statistically significant increase above basal after aerobic exercise in the present study (Figure 3A). Although cross-study comparisons require caution, these findings raise the possibility that, in individuals with obesity, the impairment in muscle anabolic responsiveness may be more pronounced following aerobic than resistance exercise.

One plausible mechanism for the attenuated anabolic response is the persistence of exercise-induced energetic stress during the early postexercise period. Acute aerobic exercise increases ATP demand and activates AMP-activated protein kinase (AMPK) (41), a key cellular energy sensor that negatively regulates skeletal muscle protein synthesis (42). Increased AMPK activation and suppressed anabolic signaling have been observed during early recovery (e.g., ∼1 h postexercise) following resistance exercise in humans (43). The magnitude of AMPK activation also depends on exercise mode, with greater activation observed following aerobic than resistance exercise (44, 45). Moreover, exercise-induced AMPK activation may be greater and more prolonged in untrained individuals, such as the participants in the present study, than in trained or physically active individuals (46). Accordingly, the aerobic exercise session may have imposed a pronounced energetic challenge that prolonged the activation of energy-sensing pathways and constrained anabolic responsiveness to amino acid provision during early recovery. Although AMPK signaling was not measured in the present study, increased AMPK activation in skeletal muscle is a well-established response to aerobic exercise (44, 45). Thus, the existing evidence and our findings raise the possibility that skeletal muscle responsiveness to amino acid provision may be delayed rather than absent, potentially emerging later during postexercise recovery in individuals with obesity.

A second, non-mutually exclusive mechanism is the lower circulating amino acid concentrations observed during the amino acid infusion in the EX + AA group. Compared with the AA group, the EX + AA group had lower plasma concentrations of several EAA and BCAA, including leucine. These differences are unlikely to be attributable to differences in the infused amino acid dose, because infusion rates were normalized to fat-free mass, an approach that produces comparable plasma amino acid concentrations between groups under nonexercise conditions (7, 21). Thus, the observed differences suggest that prior aerobic exercise may have altered systemic amino acid disposition and signaling, although the underlying processes cannot be determined from the present study. With specific regard to leucine, the lower plasma concentration following exercise is consistent with evidence that aerobic exercise increases leucine oxidation in humans (47), including individuals with obesity (48), and which is associated with reduced circulating leucine concentrations (49). Although leucine oxidation was not measured, increased oxidation could have contributed to the lower circulating leucine availability in the EX + AA group. Studies in older adults have demonstrated that relatively modest differences in leucine availability can affect the stimulation of muscle protein synthesis (50, 51). Mechanistically, reduced plasma leucine availability may limit activation of key components of the translational machinery, thereby constraining the muscle anabolic response after exercise (52). The positive correlation, across all study participants, between plasma leucine concentration and the change in muscle protein FSR from basal during the amino acid infusion supports the possibility that lower leucine availability may have contributed, at least in part, to the attenuated response in the EX + AA group.

From a practical perspective, these findings have important implications for the design of exercise and nutritional strategies in individuals with obesity. The potential contribution of reduced circulating amino acid availability in the exercise group suggests that strategies that preserve postexercise EAA availability, particularly leucine, may help mitigate the attenuated anabolic response. However, the amino acid infusion used in the present study produced large and sustained elevations in plasma amino acid concentrations that exceed those typically achieved under physiological feeding conditions. This observation suggests that reduced amino acid availability was unlikely to be the sole mechanism and that simply increasing dietary protein or amino acid provision may not fully restore the anabolic response. Importantly, these findings do not imply that aerobic exercise is broadly detrimental to skeletal muscle protein anabolism in individuals with obesity. Rather, they suggest that exercise/immediate postexercise recovery may not be the optimal temporal window for amino acid provision to stimulate skeletal muscle anabolism in this population. Accordingly, strategies intended to support skeletal muscle anabolism may need to consider the timing of protein or amino acid provision relative to the completion of aerobic exercise. Future studies should directly compare the effects of immediate and delayed nutrient provision on muscle anabolism in individuals with obesity.

We acknowledge that the sample size of the present study was modest. Nevertheless, the magnitude and overall consistency of the observed effects across participants support the biological relevance of the primary endpoint, muscle protein synthesis. We did not directly assess the molecular mechanisms underlying the attenuated response; therefore, the proposed mechanistic explanations remain speculative, and further studies incorporating molecular signaling measurements may help delineate the cellular and molecular pathways contributing to this effect. Importantly, although molecular measurements would provide mechanistic insight, muscle protein synthesis is itself an integrated physiological endpoint with direct relevance to skeletal muscle anabolism and muscle physiology.

In conclusion, in individuals with obesity, the increase in mixed-muscle protein synthesis elicited by increased amino acid availability is markedly attenuated during the immediate recovery from acute aerobic exercise. These findings indicate impaired skeletal muscle responsiveness to nutrient-derived anabolic stimuli during early postexercise recovery and suggest that nutrient timing may influence the anabolic response in this population. Future studies need to assess the temporal dynamics of postexercise anabolic sensitivity to determine when a delayed increase in plasma amino acid availability enhances the muscle protein synthetic response following aerobic exercise in individuals with obesity.

## Author Contributions

CSK conceived and designed the research; KAJ, EDSF, LRR, EDF, and CSK performed the experiments; KAJ, EDSF, LL, HG, MB, and CSK analyzed data; KAJ and CSK interpreted the results; KAJ and CSK prepared the figures; KAJ and CSK drafted the manuscript; KAJ, EDSF, LRR, EDF, HG, MB, and CSK edited and revised the manuscript; all authors approved the final version of the manuscript.

## Funding

The study was supported by National Institutes of Health/National Institute of Diabetes and Digestive and Kidney Diseases grants R01DK094062 and R01DK123441 (CSK).

## Disclosures

No conflicts of interest, financial or otherwise, are declared by the authors.

---

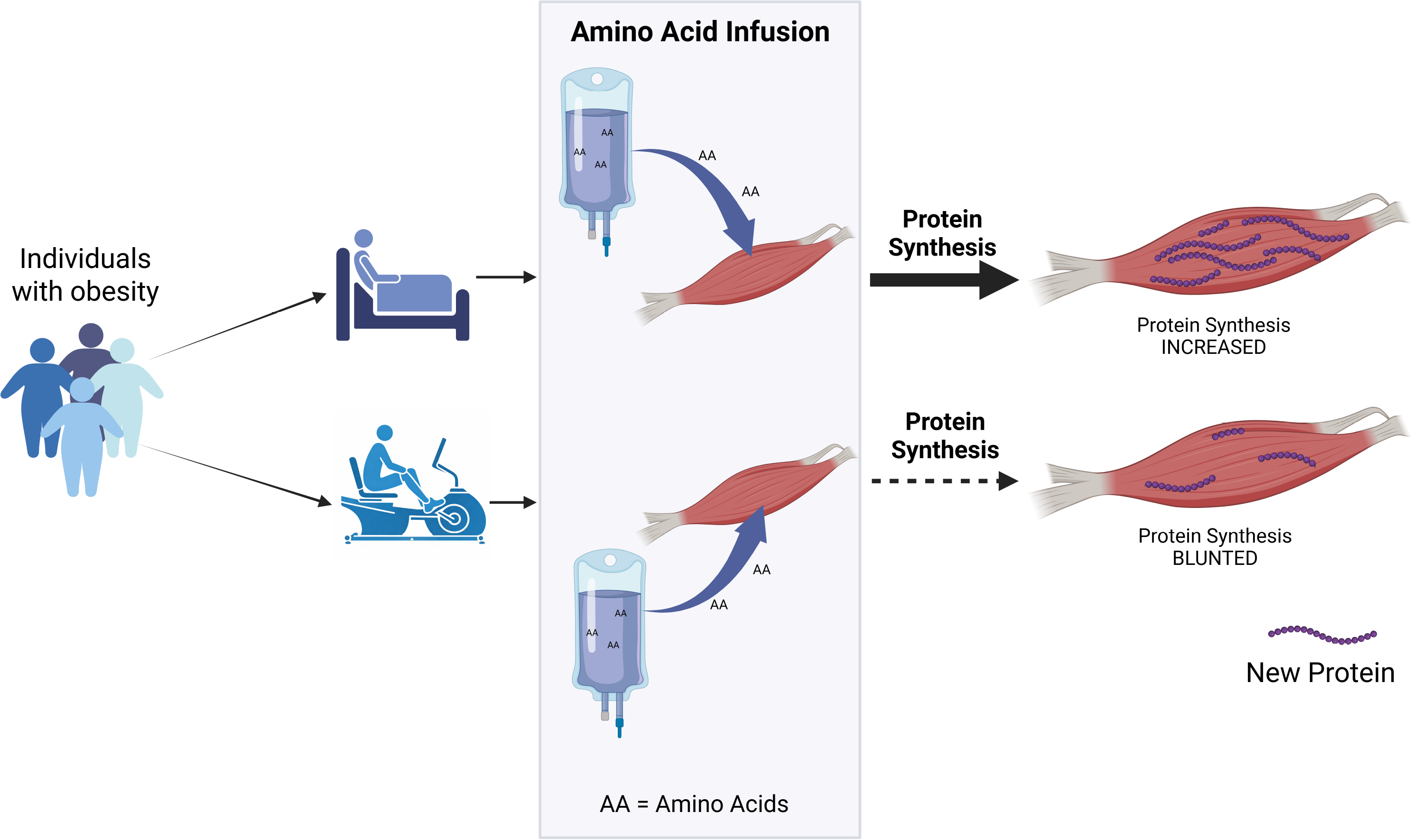

## Notes

### Competing Interest Statement

The authors have declared no competing interest.

### Summary of Updates

This version of the manuscript has been revised to include an additional author, update the title, reorganize the abstract for clarity, and add a figure depicting the experimental design.

